# Ligand Entry into Fatty Acid Binding Protein via Local Unfolding instead of Gap Widening

**DOI:** 10.1101/782664

**Authors:** T Xiao, Y Lu, J Fan, D Yang

## Abstract

Fatty acid binding proteins (FABPs) play an important role in transportation of fatty acids. Despite intensive studies, how fatty acids enter the protein cavity for binding is still controversial. Here, a gap-closed variant of human intestinal FABP was generated by mutagenesis, in which the gap is locked by a disulfide bridge. According to its structure determined here by NMR, this variant has no obvious openings as the ligand entrance and the gap cannot be widened by internal dynamics. Nevertheless, it still uptakes fatty acids and other ligands. NMR relaxation dispersion, chemical exchange saturation transfer and hydrogen-deuterium exchange experiments show that the variant exists in a major native state, two minor native-like state, and two locally unfolded states in aqueous solution. Local unfolding of either βB–βD or helix 2 can generate an opening large enough for ligands to enter the protein cavity, but only the fast local unfolding of helix 2 is relevant to the ligand entry process.

**Statement of Significance:** Fatty acid binding proteins transport fatty acids to specific organelles in the cell. To enable the transport, fatty acids must enter and leave the protein cavity. In spite of many studies, how fatty acids enter the protein cavity remains controversial. Using mutagenesis and biophysical techniques, we have resolved the disagreement and further showed that local unfolding of the second helix can generate a transient opening to allow ligands to enter the protein cavity. Since lipid binding proteins are highly conserved in 3D structures and ligand binding, all of them may use the same local unfolding mechanism for ligand uptake and release.

## Introduction

Fatty acid binding proteins (FABPs) are a family of specific carrier proteins that actively facilitate the transport of fatty acids to specific organelles in the cell for metabolism, storage, and signaling (1). They are critical meditators of metabolism and inflammatory processes and are considered as promising therapeutic targets for metabolic diseases (2). Nine types of FABPs are found in the cytosol of a variety of mammalian tissues (3). Different FABPs from human share relatively low sequence identities, but they adopt similar 3D structures with a slightly elliptical β barrel comprising two nearly orthogonal five-stranded β-sheets, a cap with two short α-helices, and a large cavity filled with water (Fig. 1A) (2, 3). Nearly all structures obtained in both crystal and solution states show no obvious openings (4), but ligands can access the binding site located inside the cavity. Knowledge of how ligands enter the cavity is important for manipulating FABP’s function by blocking the ligand entrance. Previous molecular dynamics (MD) studies suggested three possible ligand entry sites for ligand and water to enter into or exit from the protein cavity: E_I_, located in the cap region involving the second α-helix (α2) and βC-βD and βE-βF turns; E_II_, the gap between βD and βE; and E_III_, in the area around the N-terminus (5–8). Very recently, our studies on human intestinal FABP (hIFABP) have showed that opening the cap by swinging the two helices away from the barrel is dispensable for ligands to enter the cavity, and further demonstrated the existence of a minor conformational state that undergoes transient local unfolding in α2 on sub-millisecond timescale and thus provides a temporary opening in the E_I_ region for ligand entry (9). Nevertheless, we could not exclude the presence of E_II_ and E_III_.

**Figure 1.**
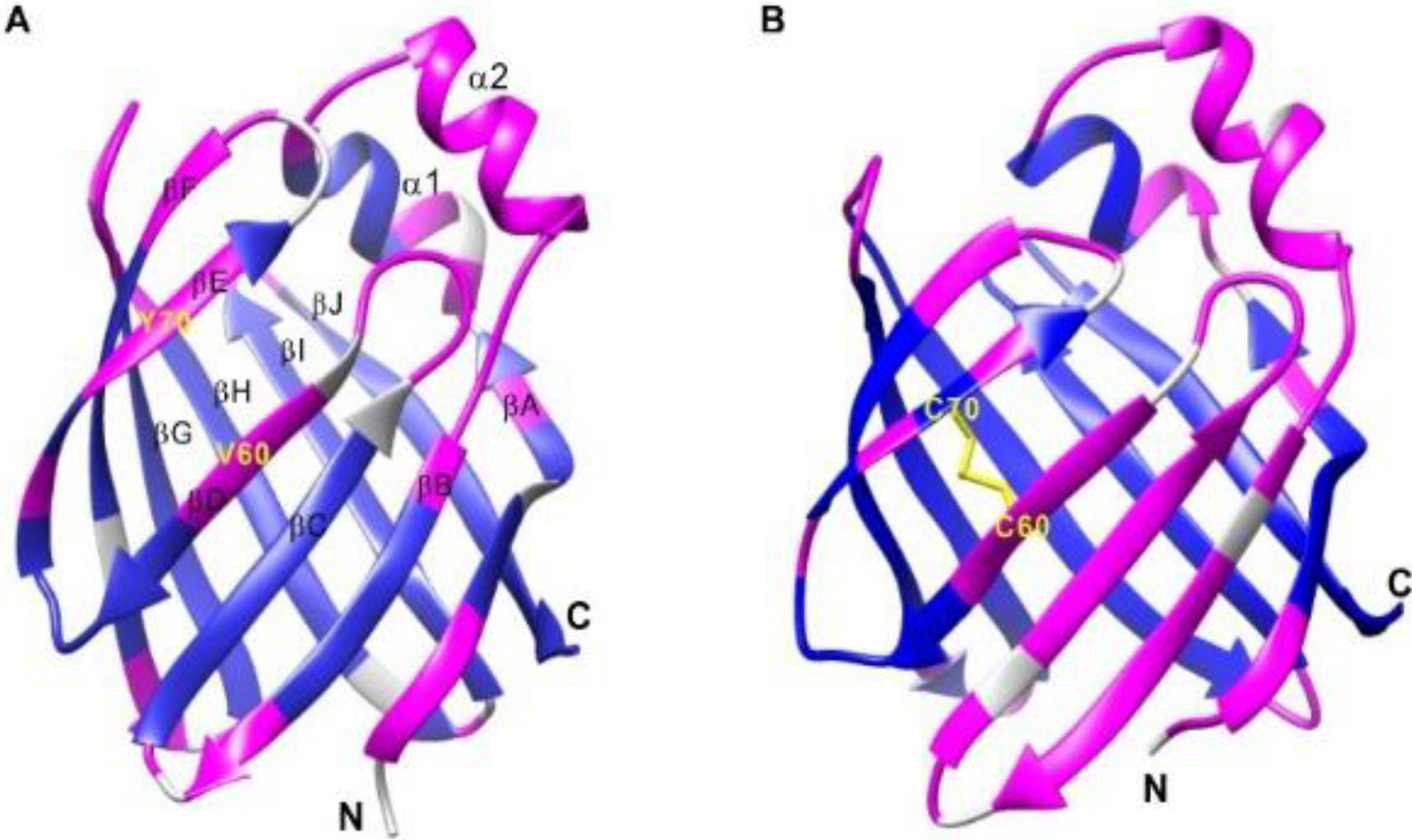
Structures of WT hIFABP (A) and its V60C/Y70C variant (B). Residues with HDX rates smaller than 0.01 s^−1^ are indicated in blue, otherwise in pink. The residues with unavailable data are in grey. The disulfide linkage is shown in yellow sticks.

There is a gap in the β-sheet of the barrel structure, where no main-chain hydrogen bonds exist between βD and βE (Fig. 1A) (10). This is common to intracellular lipid binding proteins including FABPs, suggesting that the gap may play a role in lipid uptake and release (10). Recent MD simulations on human heart FABP show that the gap between βD and βE can open to become significantly wider (5, 8). In addition, MD simulations on a human myelin protein P2, which is a member of the FABP superfamily, indicates that a large-scale opening of the barrel between βD and βE can occur through flapping βE and βF out (7). Upon this opening, the internalized cavity becomes accessible by ligands. Although computational studies suggest that opening of the barrel is likely a general mechanism for ligand entry into FABPs, experimental evidence is still lacking.

To investigate if ligands indeed enter the cavity through E_II_ via widening the gap, here we generated a gap-closed variant of hIFABP by introducing a disulfide linkage between βD and βE. Structure characterization reveals that the variant adopts a structure without obvious openings and its gap cannot be widened substantially through internal dynamics. Similar to the wild type (WT) hIFABP, the variant still takes up fatty acids and other ligands. Dynamics characterization shows that the variant exists in multiple conformational states in aqueous solution and the local unfolding process from the major native “closed” state to a minor “open” state with a locally unfolded helix 2 is relevant to ligand uptake and release.

## Materials and Methods

### Protein Sample Preparation and NMR Spectroscopy

The construct of hIFABP mutant V60C/Y70C was generated using a two-step polymerase chain reaction (PCR) scheme. The mutant (variant) was expressed in *E. coli*, purified and delipidated using the protocol described previously (11). For structure determination, a ^13^C,^15^N-labeled sample was used to acquire NMR triple resonance data including HNCA, HN(CO)CA, MQ-HCCH-TOCSY, and ^13^C,^15^N-edited NOESY (12, 13). ^15^N-labeled samples were employed for Car-Purcell-Meiboom-Gill (CPMG) and chemical exchange saturation transfer (CEST) experiments at 30 °C and hydrogen-deuterium exchange (HDX) experiments at 25 °C. Unlabeled protein samples were used for stopped-flow and affinity measurement experiments at 20 °C. All NMR experiments were performed on samples containing ~1 mM protein, 20 mM sodium phosphate, and 50 mM NaCl on a Bruker 800 MHz instrument equipped with a cryoprobe. Except the samples used for HDX at pH 7.2 and for stopped-flow at pH 9.4, other samples were at pH 7.0.

The NMR data were processed using NMRPipe (14) and analyzed for resonance assignment using Sparky. NMR resonance assignment and structure determination of the V60C/Y70C variant was achieved using a NOESY-based strategy described previously (15). On the basis of distance constraints derived from unambiguous NOE assignments and dihedral angle constraints derived from chemical shifts, the structure was calculated with Xplor-NIH (16) using the standard simulated annealing method.

HDX rates were determined from NMR signal intensity changes with HDX time after dissolving lyophilized protein into heavy water solution. Each ^1^H-^15^N HSQC was acquired with an inter-scan delay of 0.3 s and 4 scans (total experimental time of 149 s) using the so-fast HSQC scheme. Amide hydrogen exchange rates were measured in water solution using the radiation-damping-based water inversion method (17).

^15^N relaxation dispersion (RD) data were acquired with a CW decoupling and phase-cycled CPMG method (18, 19) using a constant time relaxation delay of 30 ms and inter-scan delay of 2 s. RD data at 16 different CPMG fields from 33.3 to 1000 Hz were collected by varying the separation of CMPG pulses. To estimate uncertainties in the apparent relaxation rates, the measurements at a CPMG field of 66.6 Hz were repeated three times.

CEST profiles were obtained at two different weak saturation fields (13.6 and 27.2 Hz) with a saturation time of 0.5 s and inter-scan delay of 1.5 s (20, 21). For each saturation field, 51 HSQC-based spectra were acquired using a series of ^15^N carrier frequencies ranging from 106 to 131 ppm at a spacing of 0.5 ppm. The uncertainties of the data points were estimated from the standard deviation of the points over a region far away from the CEST dips.

### RD and CEST data analysis

To extract conformational exchange parameters, RD and CEST data of the residues displaying three obvious CEST dips were simultaneously fitted to the following 4-state exchange model (Fig. 2) as described previously (22). Error estimation also followed the previous method (22).

**Figure 2.**
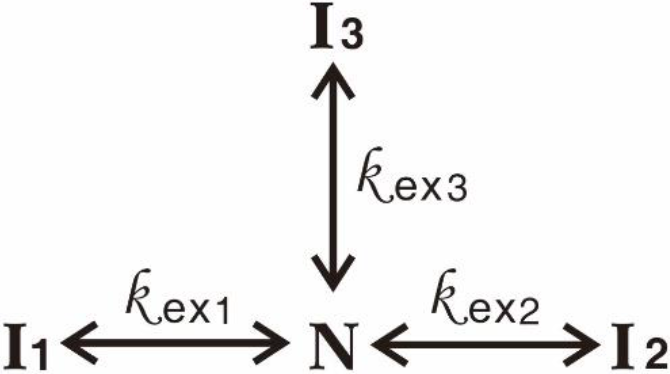
4-state exchange model. k_ex1_ (k_ex2_, k_ex3_) is the total exchange rate between state N and state I_1_ (I_2_, I_3_).

For the residues exhibiting three obvious dips, their chemical shifts in the minor states were also obtained from the global fitting mentioned above. For the residues showing significant conformational exchange contribution to transverse relaxation (R_ex_) and exhibiting one or two CEST dips, their chemical shifts in the minor states were determined by fitting the RD and CEST data of each residue to the 4-state model by fixing the exchange rates and populations of minor states at the values derived from the global fitting.

### Stopped-flow

All the stopped-flow experiments were conducted on an Applied Photophysics spectrometer by mixing protein (4 μM protein, 50 mM NaCl, pH 9.4) and oleic acid (50 mM NaCl, pH 9.4) solutions in equal volumes. At each oleic concentration, the experiment was repeated 10 times and the average data were used to extract apparent binding rates. To extract apparent binding rates, the stopped-flow traces were fitted to a bi-exponential function as described previously (9).

### ANS titration

The binding affinity of ANS to the gap-closed variant was measured using a Shimadzu RF-5301 fluorescence spectrometer. The change of ANS fluorescence intensity with protein concentration (F) was fitted to a one-site binding model:

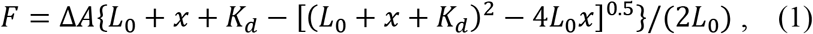

where L_0_ is the total ANS concentration (1 μM), x is the total protein concentration, K_d_ is the dissociation constant, and ΔA is the difference of fluorescence intensities between the protein-free and protein-bound ANS.

## Results and Discussion

### Structure of gap-closed hIFABP variant

A hIFABP mutant was generated by introducing a disulfide linkage between βD and βE, which was achieved by mutating V60 located in βD and Y70 in βE into cysteine (Fig. 1). The mutant displays substantially different ^1^H-^15^N NMR correlation spectra in the absence and presence of reducing agent DTT (Fig. S1), indicating that a disulfide bond exists in the mutant in the absence of reducing agents. The ^13^C_β_ chemical shifts of C60 (44.1 ppm) and C70 (45.3 ppm), which are typical for oxidized cysteine, further demonstrate the formation of a disulfide bond. To examine if introduction of the disulfide bond induces structural changes, the structure of the mutant was solved based on distance and dihedral angle restraints obtained from triple-resonance NMR experiments (Tab. S1). The mutant is very similar to the wild type (WT) protein in overall structure (Fig. 1, Fig. S2) and has no obvious openings on the protein surface (Fig. S3). Introduction of the disulfide bond reduces the gap between βD and βE by ~1 Å. Nevertheless, there are still no main-chain hydrogen bonds between βD and βE. In addition, the upper part of βB-βD of the variant orientates slightly more outward than that of the WT hIFABP (Fig. S2). Since the middle of βD is connected with the middle of βE by a covalent linkage, widening of the gap between these two strands should be insignificant (< 1 Å) by internal dynamics, if such opening can happen. Thus, the V60C/Y70C hIFABP mutant is also referred to as gap-closed hIFABP variant.

### Binding of ligands to gap-closed hIFABP variant

Except fatty acids, FABPs also bind other lipophilic molecules such as 1-anilinonaphthalene-8-sulfonic acid (ANS) that is an excellent fluorescent probe. Previous studies have shown that ANS resides in the fatty acid binding site and binds IFABP in a 1:1 ratio, the same as fatty acids (11, 23, 24). In this study, ANS was used as a fatty acid analog to test ligand binding to the gap-closed variant. On the basis of fluorescence titration, the gap-closed variant still binds ANS (Fig. S4), even though βD and βE are covalently linked and the gap cannot be widened more than 1 Å through internal dynamics. The variant with a disulfide bond has an ANS binding affinity of 10.2 ± 0.3 μM, which is similar to that for its reduced form without a disulfide linkage (11.2 ± 0.4 μM), and larger than that for the WT protein (7.1 ± 0.2 μM), suggesting that the sidechains of V60 or/and Y70 are likely involved in interactions with ANS. The results show that opening the β barrel between βD and βE is unnecessary for ligands to enter the protein cavity for binding, and imply that opening another region should occur through protein structural changes.

### Coexistence of multiple conformational states of gap-closed hIFABP

Similar to WT hIFABP, the gap-closed variant has no obvious openings. To reveal how ligands enter the cavity, we probed conformational exchanges of the variant using NMR relaxation dispersion (RD) and CEST experiments. 75 out of 131 residues displayed RD with R_ex_ values larger than 3 s^-1^ on an 800 MHz spectrometer (Fig. 3a, c, e and g), 6 residues had R_ex_ values between 2 – 3 s^-1^, 31 residues had Re_x_ values smaller than 2 s^-1^, and 18 residues with peak overlapping or weak signals were excluded for analysis. Here R_ex_ is defined as R_2_^eff^(1000) – R_2_^eff^(33), where R_2_^eff^(1000) and R_2_^eff^(33) are the relaxation rates measured at CPMG fields of 1000 and 33.3 Hz, respectively. The RD data indicate that at least one “invisible” state (I_1_) is in dynamic equilibrium with the observed native state (N) on millisecond timescale. Among the 75 residues with R_ex_ > 3 s^-1^, 18 residues each exhibited three obvious dips (Fig. 3d) that correspond to one native state and two minor states, 27 residues each displayed two obvious 2 dips (Fig. 3b, f), and 30 residues had only one dip (Fig. 3h) in their CEST profiles. The CEST data suggest the presence of at least two additional “invisible” minor states (I_2_ and I_3_) that undergo conformational exchanges with state N on sub-second timescale. To obtain structural information of the “invisible” states and kinetics parameters for conformational exchange processes, a 4-state model (Fig. 2) was used to fit both the CEST and RD data simultaneously.

**Figure 3.**
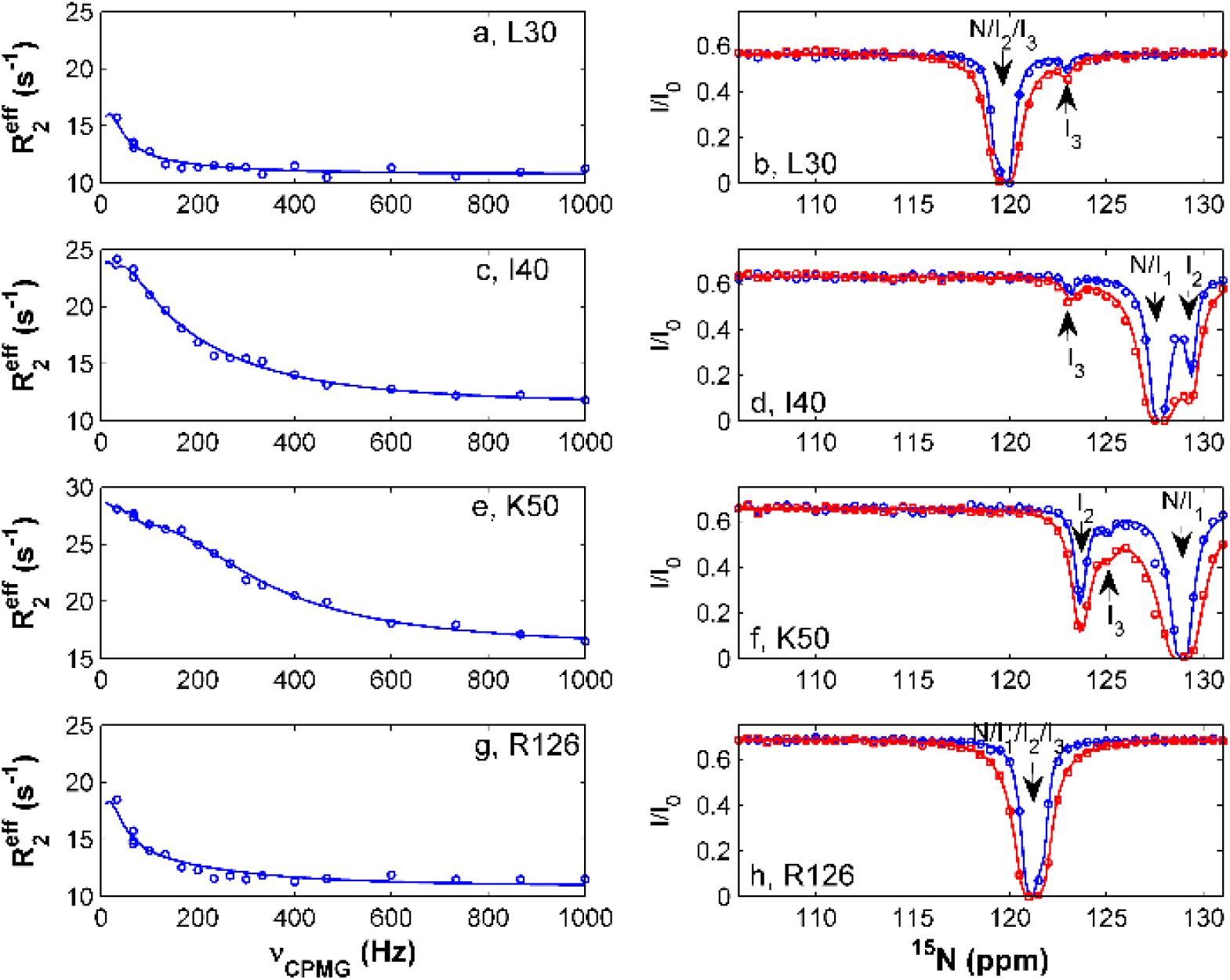
Representative RD (**a,c,e,g**) and CEST (**b,d,f,h**) profiles. The experimental CEST data at rf fields of 13.6 and 27.2 Hz are indicated by ‘o’ and ‘□’, respectively. The solid lines are best fits obtained with a 4-state model. The locations (or chemical shifts) of states N, I_1_, I_2_, and I_3_ in the CEST profiles are indicated by arrows.

Fitting the data from all the residues with three obvious dips globally, we obtained populations of states I_1_, I_2_ and I_3_ (p_I1_ = 3.4 ± 0.3%, p_I2_ = 5.3 ± 0.2%, and p_I3_ = 1.8 ± 0.9%) and their respective exchange rates with state N(k_ex1_ = 1629 ± 116 s^-1^, k_ex2_ = 82 ± 5 s^-1^, and k_ex3_ = 16 ± 8 s^-1^). ^15^N chemical shifts of the minor states determined from the data fitting are listed in Table S2.

For other residues with 1 - 2 CEST dips and R_ex_ > 2 s^-1^, their chemical shifts in minor states were obtained from data analysis by fixing k_ex1_, p_I1_, k_ex2_, p_I2_, k_ex3_, p_I3_ at the values derived from the global fitting, which are also listed in Table S2.

Folded and unfolded proteins have very distinct ^15^N chemical shifts. Comparing the chemical shifts of the minor and native states (Tab. S2), we can see that states I_1_ and I_2_ are much more similar to state N than unfolded state U (Fig. 4a-d), but state I_3_ is significantly different from both states N and U. For state I_3_, residues located in the N-terminal region of βA (F2–W6), βB–βD (N35–L64), and C-terminal end of α2 (L30–D34) are similar to the unfolded state, while residues located in other regions are similar to the native state in terms of ^15^N chemical shift (Fig. 4e-f). For instance, the chemical shifts of residues K37–E43 in state I_3_ each differ from those in state N by more than 3.4 ppm, but deviate from those in state U by less than ~1.0 ppm (Tab. S2, Fig. 4e-f). Therefore, states I_1_ and I_2_ are native-like, while state I_3_ is partially unfolded in which the N-terminal region of βA, βB–βD, and C-terminal end of α2 are mainly disordered. The unfolding rate from state N to state I_3_ is about 0.3 s^-1^ (p_I3_*k_ex3_), significantly smaller than the conversion rates from state N to I_1_ (~55 s^-1^) and I_2_ (4 s^-1^).

**Figure 4.**
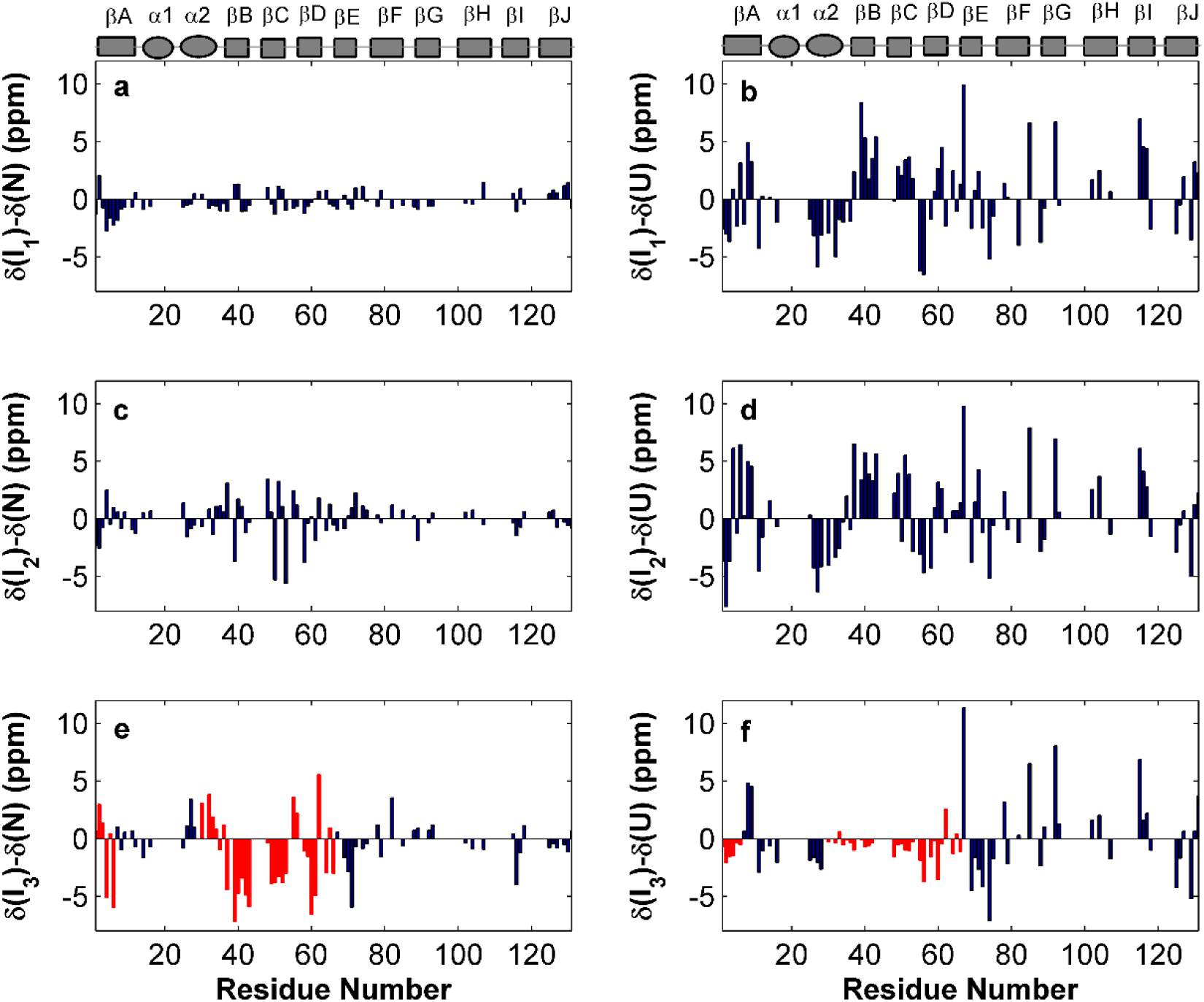
Differences of ^15^N chemical shifts between states I_1_ and N (a), between state I_1_ and unfolded state (U) (b), between states I_2_ and N (c), between states I_2_ and U (d), between states I_3_ and N (e), and between states I_3_ and U (f). In (e) and (f), the residues in the N-terminal region of βA, C-terminal region of α2, and βB - βD are marked in red. Secondary structure elements are shown on the tops of (a) and (b).

To examine if there are other conformational exchanges in the gap-closed variant, amide hydrogen exchange experiments were performed. 62 out of 131 residues displayed ^1^H-^15^N correlations in the first HDX spectrum which was recorded with a total acquisition time of 149 seconds and a dead time of ~160 seconds. Their HDX rates are listed in Table S3. For other residues with HDX rates larger than 0.2 s^-1^, the exchange rates of their backbone amides with water hydrogen were measured (Tab. S3). As expected, the residues not involved in H-bonding have large amide hydrogen exchange rates and small protection factors (PFs). Interestingly, all residues located in α2, βB and βC of the gap-closed variant have small PFs (<100) (Tab. S3, Fig. 1) even though their backbone amides are involved in H-bonding. In contrast, most residues located in βB and βC of the WT hIFABP and its cap-closed variant have very large protection factors (>1000) (9, 22). The exchange rates for most residues at pH 7.2 were 1.4 – 2.0 times as large as those measured at pH 7.0 (Tab. S3), indicating that the amide hydrogen exchange can be described by the EX2 model (25). Using this model, the population of an amide in a disordered form can be approximated as 1/PF. According to the PFs of the amides involved in H-bonding (Tab. S3), the populations of the disordered form were ~15-30% for α2, ~2% for N-terminal region of βA, ~0.5-8% for βB and βC, and <0.1% for α1 and βF–βJ. It is noteworthy that the populations estimated from the PFs are error-prone, strongly depending on the predicted exchange rates. This result further supports that states I_1_ (p_I1_ = 3.4%) and I_2_ (p_I2_ = 5.3%) are native-like rather than unfolded, while state I_3_ (p_I3_ = 1.8%) is partially unfolded in which only βB, βC, and α2 are mainly disordered.

The population of the disordered form for α2 estimated from PFs (~15-30%) is much larger than the population of state I_3_ derived from our CEST data (1.8%), suggesting the presence of an additional state in which only α2 is disordered. This locally unfolded state is denoted as I_4_. Since state I_4_ was not observed by CEST and RD experiments, its exchange rate with state N should be significantly smaller than k_ex3_ (16 s^-1^) or much larger than k_ex1_ (1600 s^-1^). In the former case (slow exchange regime), the residues in α2 should give rise to two sets of ^1^H-^15^N correlation peaks corresponding to states N and locally unfolded I_4_. In fact, only one set of peaks corresponding to state N were observed, indicating that state I_4_ must undergo fast exchange with state N. The rate of the fast exchange should be on the microsecond timescale so that the exchange could not be detected by ^15^N CPMG experiments.

### Functionally relevant conformational exchange

The partially unfolded state I_3_ should have an opening large enough for ligands to enter the protein cavity. To examine if this partial unfolding process is relevant to the uptake of fatty acids, we measured the apparent association rate of oleic acid binding to the gap-closed hIFABP using fluorescence stopped-flow. The stopped-flow data could be fitted well to a bi-exponential function instead of mono-exponential function when oleic acid concentrations were lower than 25 μM (Fig. S5). This suggests that the ligand binds the protein in at least two steps. At higher oleic acid concentrations, the intrinsic protein fluorescence signal started to decay after a mixing time of ~4 ms (Fig. S5) due to the quenching effect, which was not observed for the wild type protein (9). In this case, only the apparent association rate for the fast step could be estimated. The fast apparent rates increased with oleic acid concentrations initially, and then gradually reached a plateau with further increase of ligand concentration (Fig. 5). This result suggests that there is a rate-limiting step before the ligand association step and the rate limit is ~1000 s^-1^ for the gap-closed variant. On the other hand, the rates for the second association step were small (~ 20 s^-1^) and nearly independent of ligand concentrations.

**Figure 5.**
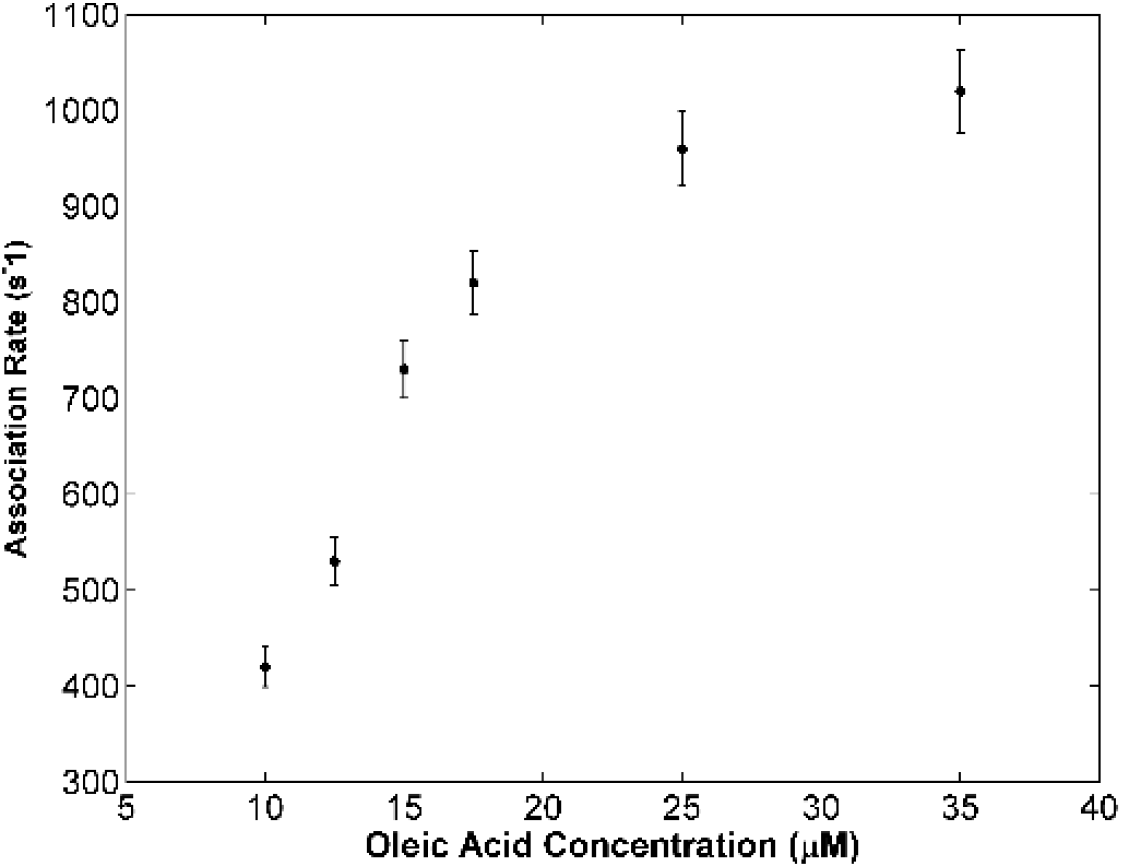
Dependences of apparent association rates for the fast step on ligand concentrations.

The binding kinetics results for the gap-closed mutant are similar to those for the cap-closed mutant and wild type protein (9). Therefore, the three-step binding model for the cap-closed variant described previously should be applicable to this gap-closed variant too. In this model, the maximal apparent association rate for the fast step should be smaller than the conversion rate from a closed state to an open state since the protein stays mainly in a closed state. If fact, the maximal apparent association rate (~1000 s^-1^) is much larger than the conversion rates from state N to I_1_, I_2_ and I_3_ (<55 s^-1^). So the local unfolding process from state N to I_3_ and other two conformational exchange processes revealed by RD and CEST are irrelevant to the ligand entry for the gap-closed variant. Different from state I_3_, state I_4_ with a disordered α2 is in fast exchange with state N, which is similar to the locally unfolded state of the cap-closed variant. For WT hIFABP and its gap-closed and cap-closed variants, the PF values are smaller than 100 for R28-A32 in α2 but larger than 1000 for V17-M21 in α1 (9, 22) even though the amides of these residues are all involved in H-bonding and have similar water accessibility, suggesting the presence of a minor state with locally unfolded α2 for all hIFABP proteins. Our recent study (9) on the cap-closed mutant showed that local unfolding of α2 is the rate-limiting step for ligands to enter hIFABP cavity for binding. Therefore, it is reasonable to assume that the unfolding of α2 also controls the entry of ligands into the gap-closed variant and the conversion rate from state N to I_4_ is close to the maximal ligand association rate (~1000 s^-1^).

## Conclusions

In summary, we have generated a gap-closed hIFABP variant with a disulfide linkage between strands βD and βE, which has a similar 3D structure to the WT protein and has no obvious opening on its surface. The variant always stays in a gap-closed state due to the presence of the disulfide linkage, but it still takes up fatty acids and other lipophilic ligands. Thus, ligands do not enter the protein cavity for binding via opening the gap, rejecting the previously proposed gap-opening mechanism. The native state of the gap-closed variant is in dynamic equilibrium with two minor native-like states (I_1_ and I_2_), one locally unfolded state in which the N-terminal region of βA, βB–βD, and α2 are mainly disordered (I_3_), and one locally unfolded state in which α2 is mainly disordered (I_4_). The conversion rate from state N to I_3_ (0.3 s^-1^) is much smaller than the apparent association rate of oleic acid, indicating that ligands do not enter the protein cavity through the opening created by local unfolding of the βB–βD region. Local unfolding of α2 is fast and thus relevant to the ligand entry process.

## Author contributions

D. Yang designed the research. T. Xiao and Y. Lu performed the experiments and analyzed the data. J. Fan analyzed the NOESY data and calculated the structure. Y. Lu and D. Yang wrote the manuscript.

## Acknowledgements

This work was supported by Singapore Ministry of Education Academic Research Fund Tier 2 (MOE2017-T2-1-125).

